# PyRID: A Brownian dynamics simulator for reacting and interacting particles written in Python

**DOI:** 10.1101/2024.11.26.625395

**Authors:** Moritz Becker, Nahid Safari, Christian Tetzlaff

**Affiliations:** Group of Computational Synaptic Physiology, Department of Neuro- and Sensory Physiology, University Medical Center Göttingen; III. Physical Institute - Biophysics, Faculty of Physics, University of Göttingen

## Abstract

Recent technological developments in molecular biology led to large data sets providing new insights into the molecular organisation of cells. To fully exploit their potential, these developments have to be complemented by computer simulations that allow to gain in-depth understanding on molecular principles. We developed the Python-based, reaction-diffusion simulator PyRID, integrating many features for the efficient simulation of molecular biological systems. Amongst others, PyRID is capable of simulating unimolecular and bimolecular reactions as well as pair-interactions to assess the dynamics resulting from individual interacting proteins to polydisperse substrates consisting of many different molecules. Furthermore, PyRID supports mesh-based compartments and surface diffusion of particles, enabling analyses of the interaction between (trans-)membrane proteins with intra- and extracellular proteins. PyRID is written entirely in Python, which is a programming language being known for its readability and easy accessibility, such that the scientific community can easily extend PyRID to its current and future needs.

## 1 Introduction

Cells are intricate structures comprised of biomolecules interacting continuously to regulate cellular functions. Recent advancements in experimental techniques provide insights into the distribution, movements, and reactions of molecules within cells. However, observing the spatiotemporal interplay of many different molecule species is, despite the great advances in technologies such as super-resolution microscopy or mass spectrometry, beyond reach. Computational modeling is essential for understanding molecular localization, movement, interactions, and to derive the underlying driving principles. Hereby molecular dynamics simulations are well suited to integrate information of single molecules and to offer insights into the resulting dynamics of populations of molecules. Therefore, they are a great tool that is used in different disciplines of physics, material sciences, or chemistry. However, investigating molecular processes in biological systems implies special demands on the simulators: The intracellular as well as the extracellular fluid is in general highly polydisperse, meaning that it consists of many different molecule or protein species [1, 2]. Proteins take up most of the intracellular space [3, 4], making a biological cell a dense molecular environment. The cell membrane as well as organelles shape the space of molecule movement [5, 6], transmembrane proteins like ion channels move onto the surface of the membrane, interacting with proteins that are in the intracellular fluid [7, 8]. To simulate physical or chemical systems, often the thermodynamic assumption of an isolated system is being used to describe the system’s environment. In a biological cell, each part continuously interchanges ions, proteins, etc. with its surroundings, making the formulation of an isolated system difficult, which has to be considered in the simulations. Over the last years, multiple simulators have been developed to adhere biological requirements. Based on these simulators, we have developed PyRID (Python Reaction Interaction Diffusion Simulator), a Python-based, easy-to-expand simulator that is especially developed and includes many required features to simulate the molecular dynamics within biological systems such as neurons.

PyRID is a reaction-diffusion simulator that is completely written in the Python programming language and draws inspiration from other simulators like MCell, Smoldyn, and ReaDDy. PyRID is capable of simulating unimolecular and bimolecular reactions, pair-interactions, mesh-based compartments, and surface diffusion. While the aforementioned simulators exhibit impressive performance, in the following paragraph, we will discuss their strengths and weaknesses to elucidate the rationale behind PyRID’s development.

MCell is a stochastic simulation software that accurately models the behavior of particles in a system characterized by diffusion and non-interacting processes [9]. Non-interacting processes means that MCell is limited in simulating systems with strong molecular interactions and in accounting for molecular crowding effects and geometric restrictions. Anisotropic translational diffusion is not supported and molecule structure is only indirectly accounted for by internal state variables of the point particles. Despite these limitations, MCell provides excellent accuracy in modeling reactions and supports the use of triangulated meshes to represent cell surfaces and define the simulation space. It provides periodic, repulsive,and fixed concentration boundary conditions and includes a Python interface and BioNetGen reaction language (BNGL) support.

Smoldyn is a particle based reaction diffusion simulation software very similar to MCell [10]. Compared to MCell, Smoldyn provides in addition an approximation to the excluded volume for spheres and anisotropic translational diffusion. Due to the lack of support for energy potential functions, particle interactions by means of bonds cannot be simulated.

ReaDDy is a particle-based reaction-diffusion simulator that models the interactions and reactions of collections of particles, bridging the gap between highly detailed molecular dynamics and large-scale reaction-kinetics simulations in the spirit of MCell and Smoldyn [11]. ReaDDy uses external potentials to confine particles and supports periodic and repulsive boundary conditions. Complex multi-particle structures can be defined by topology graphs and a wide range of interaction potentials. However, unlike MCell, ReaDDy does not support 3D meshes for modeling geometrically complex compartments and surfaces. Other desirable but missing features include rigid bead models for molecule representation, anisotropic diffusion and fixed concentration boundary conditions. Also, polydispersity can cause significant performance drops.

In order to have a simulator integrating a wide spectrum of features desired for investigating biological systems, we have developed PyRID that incorporates various tools from other simulators. In addition, as the rapid development in the research of biological systems implies the need to easy adapt the source code of a simulator by individual researchers, different to the simulators discussed above, PyRID is written entirely in Python, which is a programming language being known for its readability and easy accessibility. As core component, in PyRID, a molecule can be described by one or more rigidly connected beads of arbitrary spatial configuration with corresponding anisotropic translational and rotational diffusion. On the surface of each bead, several, specific patches (represented as particles) can be defined that determine the interaction of the molecule with others, allowing multivalent protein-protein interactions. As each molecule can have a structure that is defined by beads of different sizes, using a hierarchical grid data structure, PyRID supports an efficient simulation of polydisperse systems. Triangulated mesh geometries can be formulated for examining the influence of different cell geometries on molecular organization. Furthermore, the movement of molecules can be constraint onto these geometries enabling the investigation of transmembrane protein dynamics including their interaction with intra- or extracellular proteins. Similar to MCell, PyRID allows the consideration of boundary conditions with fixed concentrations of molecules to emulate the behavior of the simulated system as compartment being in a continuous interchange with the rest of the cell. PyRID also exhibits high accuracy in simulating various types of reactions, and is readily modifiable and expandable by Python programmers. Moreover, PyRID employs Numba jit compilation to achieve efficient performance. The complete PyRID code, along with extensive online documentation and examples, is available on the GitHub repository: Code: https://github.com/MoritzB90/PyRID; Documentation: https://moritzb90.github.io/PyRID_doc/.

In the following section we will introduce the main features and rationale of PyRID. Then, we will provide different examples from computational chemistry and biology for validation and show PyRID’s performance together with some use-cases.

## 2 Theory and Methods

### 2.1 Single molecule definition

Molecules and proteins exhibit anisotropic and multivalent interactions that cannot be accurately described by isotropic energy potentials or by point-like particles [12, 13]. Although all-atom molecular dynamics simulations can accurately capture protein-protein interactions, simulating systems with numerous molecules within a reasonable time frame for processes such as protein assembly, is computationally unfeasible with current computers and algorithms. Therefore, PyRID employs coarse-graining methods [14] to represent molecular structures. Rigid bead models offer advantages in coarse-grained modeling by replacing strong and short-range interactions between atoms with a rigid bead topology, allowing for larger integration time steps. The beads do not necessarily represent single atoms or molecules, but rather the geometry of the molecule of interest, resulting in reduced computational cost. Additionally, diffusion tensors can be used to describe molecular diffusion. Multivalent protein-protein interactions can be represented by patches on the bead model surface. Please note, the accuracy of coarse-grained models depends on the choice of interaction potentials and model parameters that have to be estimated from experiments and theory. Different molecule types or species each with its own structure, volume, interaction sites, interaction types (number, arrangement, and properties of beads and patches) can be defined.

#### Translational and rotational motion of molecules

The motion of an isolated rigid bead molecule *j* in solution can be described in terms of the coupled Langevin equations for translational and rotational motion (Eqs.1, 2). We do not account for the hydrodynamic interaction between molecules as this is computationally very expensive (*O*(*N* ^2^) *− O*(*N* ^3^)) [15, 16]. For more details please refer to the following papers [17, 18, 19]. In the following, for better readability, we will omit the time-dependency of variables.

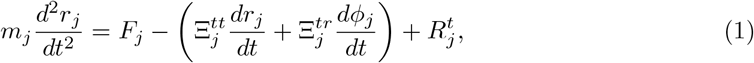

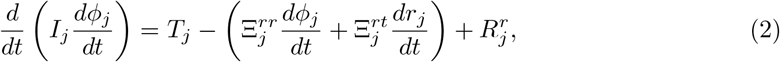

where *r*_*j*_ is the position of the center of molecule *j* and *ϕ*_*j*_ its rotation angle, *m*_*j*_ is mass and *I*_*j*_ inertia. *F*_*j*_ is the total force exerted on the molecule and *T*_*j*_ is the torque. *R*^*t*^*j* and *R*^*r*^*j* describe the random, erratic movement of the molecule due to collisions with the solvent molecules where

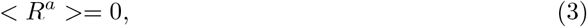

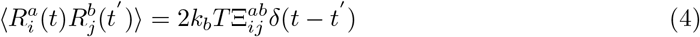

with *a, b ∈*[*t, r*], *k*_*b*_ being the Boltzmann constant, and *T* the temperature. *<*. *>* indicates the time average. Here Ξ^*tt*^, Ξ^*rr*^, Ξ^*tr*^, Ξ^*rt*^ are the translational, rotational and translation-rotation coupling friction tensors of the rigid body in the lab frame with *i* and *j* being their components. These tensors characterize the resistance encountered by the molecule due to its interactions with the surrounding medium. Additionally, the Einstein relation Ξ^*ab*^ = *k*_*B*_*T* (*D*^*ab*^)^*−*1^ relates the friction experienced by the molecule to its diffusion properties, providing insight into how the molecule’s mobility is affected by its interactions with the solvent, where *D*^*ab*^ represents the diffusion tensor. When considering low-mass particles such as molecules over extended time intervals, the acceleration of the particles may be disregarded in the portrayal of the diffusion process. Thus, we utilize overdamped Langevin dynamics, also known as Brownian motion, to describe the motion of molecules. 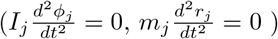

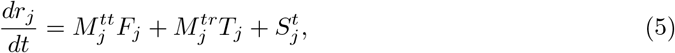

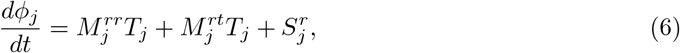

with

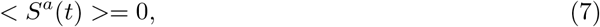

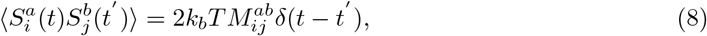

where *M* ^*tt*^, *M* ^*rr*^, *M* ^*tr*^, *M* ^*rt*^ are the translational, rotational and translation-rotation coupling mobility tensors of the rigid body in the lab frame and 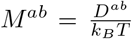 also *M* ^*rt*^ = (*M* ^*tr*^)^T^. In most cases, the influence of the translation-rotation coupling on molecular dynamics is negligible. However, translation-rotation coupling increases the complexity of the translation and rotation vector propagation algorithms. Therefore, we consider translation and rotation as being independent.

One problem with the rotational equation of motion is that several issues arise depending on how rotations are represented. For instance, propagating the rotation in terms of Euler angles will result in numerical drift and singularities [20, 21]. Therefore, especially in computer graphics, it is standard to represent rotations in unit quaternions, which is much more stable and has fewer issues in general. We also use quaternions in PyRID. An algorithm for the rotation propagator based on quaternions can be found in [22] and the PyRID documentation.

#### Mobility tensor for rigid bead models forming a molecule

In order to simulate the motion of molecules, we need to calculate the molecule’s diffusion tensor. Diffusion tensors can be also estimated experimentally [23] or using molecular dynamics simulations [24]. However, for many proteins, the diffusion tensor is unknown. Therefore, it is often more convenient to calculate the diffusion tensor directly from the coarse-grained representation of a molecule in terms of the rigid bead model. For more details, see [25] and [26]. In general, the mobility and/or diffusion tensor of an anisotropic rigid body can be calculated from the inverse of the rigid body’s friction supermatrix [27]:

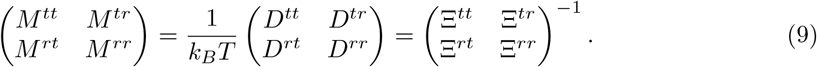

The main challenge lies in deriving an expression for the translational, rotational and translation-rotation coupling tensors of the friction supermatrix. PyRID uses a method based on a modified Oseen tensor [26, 27] to account for the hydrodynamic interaction between the beads of a rigid bead molecule in a first order approximation.

### 2.2 Particle-particle interactions

PyRID can handle external potentials and pairwise, short-ranged interaction potentials.

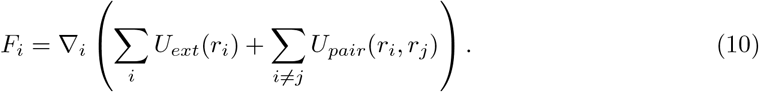

*U*_*ext*_*(r*_*i*_) represents the external potential acting on particle *i* at position *r*_*i*_, while *U*_*pair*_(*r*_*i*_, *r*_*j*_) denotes the pairwise interaction potential between particles *i* and *j* separated by a distance *r*_*ij*_. PyRID has built-in functions for often used pairwise interaction potentials, such as weak piecewise harmonic potential [11], harmonic repulsion potential [11], Continuous Square-Well (CSW) potential [28], and Pseudo Hard Sphere (PHS) potential [29]. However, PyRID does not support long-range interaction potentials including Ewald summation, and the interaction potentials have to have a cutoff distance. Nevertheless, researchers can conveniently add any short-ranged, pairwise interaction potential using Python.

### 2.3 Compartments

Cellular processes are heavily influenced by compartmentalization, and it is desirable to confine diffusion and reactions to specific regions of interest. Various approaches can be taken to restrict the movement of particles within a confined region. One method involves placing particles along the boundary of the compartment, where they can interact with particles within the compartment via a repulsive interaction potential. Another approach would be to add external potentials/force fields that restrict the motion of particles either to the volume or the surface of a compartment. This method is used in ReaDDy [11]. However, complex geometries/compartment shapes are more difficult to establish. A third approach represents the compartment geometry by triangulated meshes as is done in MCell [9]. This approach enables a separation of the simulation volume, e.g., into an extracellular and intracellular space. Accordingly, PyRID defines a compartment as a triangulated manifold mesh, which is efficiently stored in a shared vertex data structure [30] that includes an array of vertex positions and an array that stores triangle indices using three vertices (Figure 1 A).

**Figure 1:**
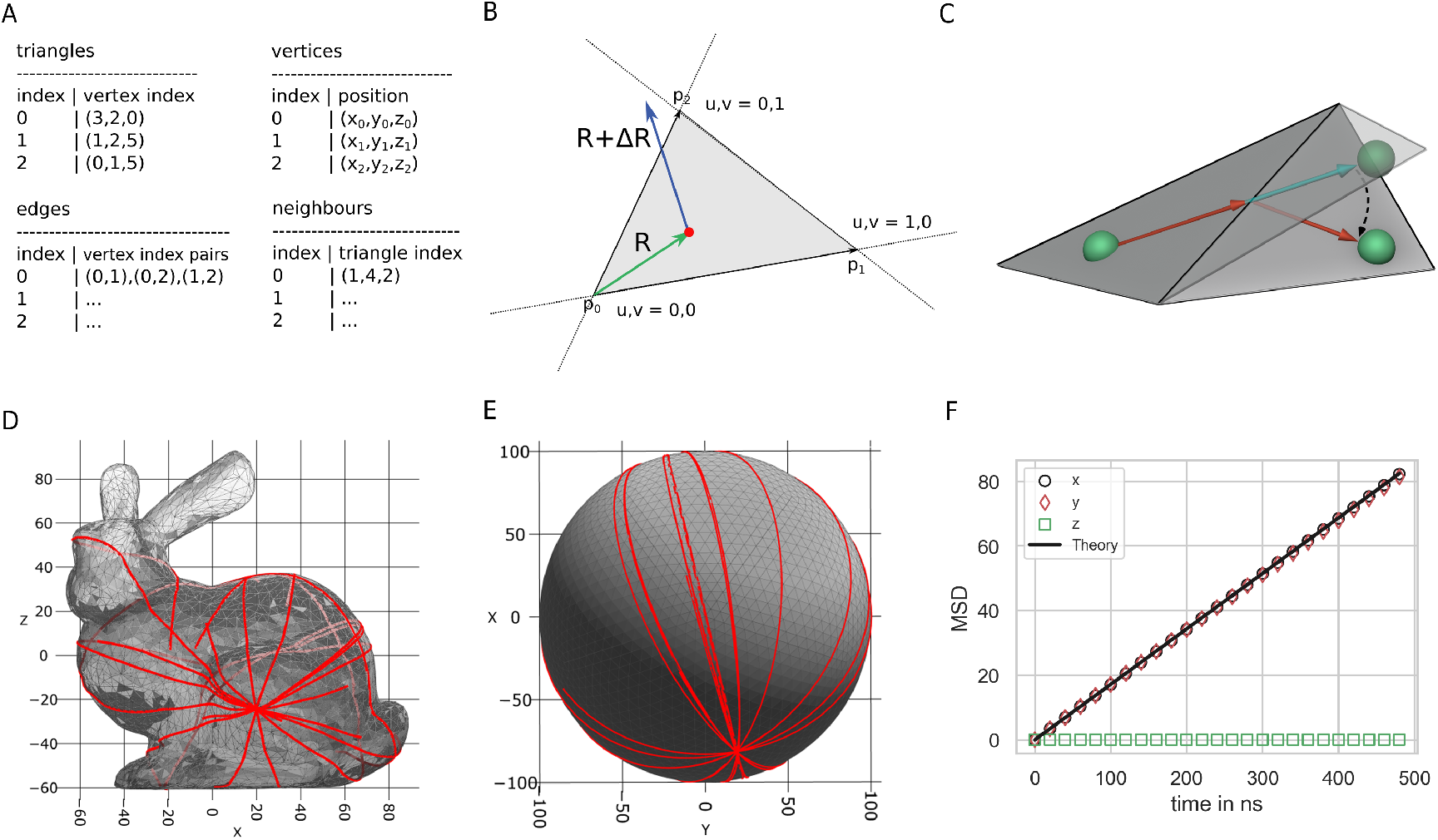
Mesh compartments and surface molecules. (A) PyRID uses triangulated meshes to represent compartments. These are kept in a shared vertex mesh data structure [30]. In addition, for neighbour search, two arrays that hold for each triangle the vertex indices of the three triangle edges and the triangle indices of the three triangle neighbours, are used. (B) Triangle vertices are ordered counterclockwise, as are edges. Efficient algorithms based on barycentric triangle coordinates are used to check whether a point lies within a triangle or whether a displacement vector intersects a triangle edge. (C) Visualization of mesh surface ray marching. If a molecule (green sphere) crosses a triangle edge, its displacement vector is advanced to the corresponding edge and then rotated into the plane of the neighboring triangle. (D,E) By the ray marching method described in the text, molecules follow a geodesic paths on the mesh surface. (F) The mean squared displacement of diffusing surface molecules is in agreement with theory, which is in 2 dimensions *MSD* = 4*Dt* (here *D* = 43 *nm*^2^*/μs*).

#### Volume molecules

In PyRID, two distinct methods are utilized to calculate the collision response of a molecule with a mesh. For large rigid bead molecules, collision detection involves neighbor searching and contact resolution, wherein each mesh triangle exerts a repulsive force on individual beads [31]. The accuracy and complexity of collision detection and force calculation depend on factors such as the type of geometries involved, surface complexity, and whether frictional forces must be considered. PyRID uses the “Point to Triangle” algorithm by [32] to calculate the distance between beads and triangles. For small, isotropic molecules or atoms, contact force calculation using repulsive forces requires a very small integration time step. To avoid this, PyRID utilizes ray tracing for collision detection and resolution, which works independently of integration time step (Figure 1 B). This approach, similar to that used in MCell [9], involves tracing the displacement vector of the molecule Δ*R* through the simulation volume and resolving collisions with the compartment boundary (mesh) via reflection. Collision tests are done using the “Fast Voxel Traversal Algorithm for Ray Tracing” introduced by [33].

#### Surface molecules

In PyRID, the lateral diffusion of surface molecules within the mesh surface is modeled with the aim of simulating the movement of transmembrane molecules such as receptors (Figure 1 C). A method similar to that used in MCell [9] is employed, wherein a molecule diffuses within the plane of a triangle until it encounters an edge. At this point, the molecule’s displacement vector Δ*R* is advanced to the edge and then rotated into the plane of the neighboring triangle, with the rotation axis determined by the shared edge. The molecule will then move in a straight line on the mesh surface, equivalent to unfolding the triangles over the shared edge to place them in a common tangent space, advancing the position vector, and folding/rotating back. The latter view provides intuitive insight into the straight-line motion of the molecule on the mesh surface using this approach.

The method sketched above is known as “Surface Ray Marching”, which we introduce briefly in the following. To effectively detect if a triangle edge has been crossed and to identify its neighboring triangle, additional data is stored alongside triangle and vertex information. Specifically, for each triangle, an array holds the vertex indices of its three edges, sorted in a counterclockwise manner, along with a separate array containing the indices of the corresponding neighboring triangles for swift reference (Figure 1 A). The triangle edge intersection test is made efficient using barycentric coordinates, where the center of the molecule (*R*_0_) and its displacement vector (Δ*R*) are described in terms of these coordinates. Similar to [34], the edges are sorted counterclockwise, starting from the triangle’s origin, simplifying the edge intersection test. To determine which edge is intersected first, the smallest distance to the respective edge is checked. The process involves advancing *R* by *R* = *R*_0_ + *t*_*i*_Δ*R* with *t*_*i*_ being the distance to the intersecting edge, adjusting Δ*R* accordingly, and transforming *R* into the local coordinate frame of the neighboring triangle. Additionally, as PyRID supports anisotropic rigid bead molecules, the molecule’s orientation needs updating for each crossed triangle. This is achieved by rotating the molecule using a rotation quaternion, with the quaternion propagation occurring through multiplication. The process terminates when *R*_0_ + Δ*R* lies within the triangle the molecule is currently located on.

### 2.4 Boundary Conditions

PyRID supports three types of boundary conditions: periodic, repulsive, and fixed concentration. Repulsive boundary conditions are implemented either through a repulsive interaction potential or via ray tracing, also see section “Volume molecules”. When using periodic boundary condition, the minimal image convention is applied to ensure that each particle interacts only with the closest image of the other particles in the system (Figure 2 B). However, it is important to note that the size of the simulation box should not be too small, as this can result in particles interacting with themselves. Since periodic and repulsive boundary conditions are common in many simulation software, we will focus on introducing the less common feature of PyRID, which is the fixed concentration boundary condition.

**Figure 2:**
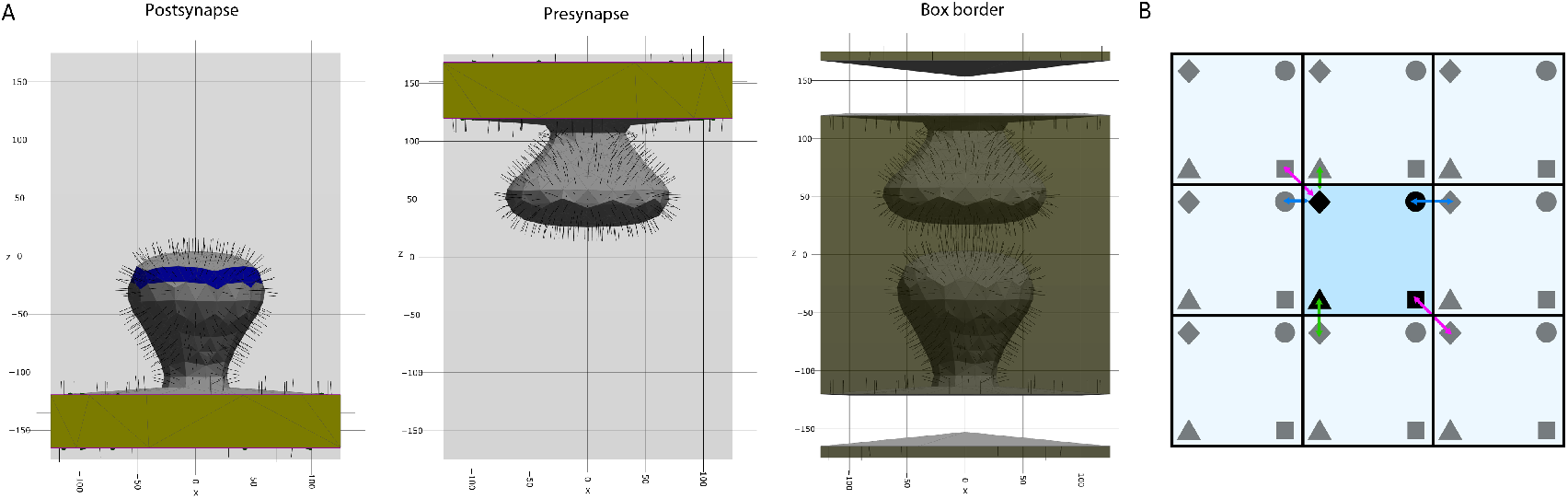
Boundaries. (A) *left, middle:* In PyRID, the user can define different face groups. Face groups can be used, e.g., to distribute molecules on specific regions of the mesh surface (blue). When a compartment intersects with the simulation box, the intersecting triangles are assigned to a transparent class (yellow), as are the corresponding edges that intersect with the boundary (purple lines). If boundary conditions are set to “fixed concentration” transparent triangles and edges act as absorbing boundaries but in addition release new molecules into the simulation volume. *right:* If mesh compartments intersect the boundary of the simulation box, the remaining part of the box boundary must also be represented by a triangulated mesh. (B) For periodic boundary conditions, PyRID follows the minimal image convention, i.e. a particle (black marker) only interacts (colored arrows) with the closest image (grey marker) of the other particles in the system.

#### Fixed concentration boundary conditions

The fixed concentration boundary condition in PyRID allows for coupling the simulation box to a particle bath, which enables simulation of a sub-region within a larger system without the need to simulate the dynamics of the molecules outside the simulation box directly (Figure 2 A). Virtual molecules outside the simulation box only become part of the simulation if they cross the simulation box boundary. In PyRID, it is possible to have mesh compartments intersect with the simulation box boundary, and molecules can enter and exit the simulation across the intersection surface or intersection curve in the case of surface molecules. The expected number of hits (*N*_*hit*_) between a molecule type and simulation box boundaries is calculated at each iteration, based on the fixed outside concentration of the molecule *C*, the diffusion coefficient *D*, and the total boundary surface area *A*. The average number of volume molecules that hit a boundary of area *A* from one side within a time step

Δ*t* can be calculated from the molecule concentration *C* and the average distance a diffusing molecule travels normal to a plane *l*_*n*_ within Δ*t* [9]:

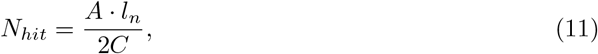

where

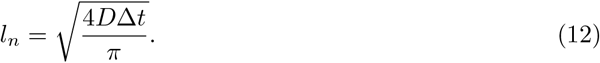

Here 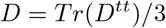 is the scalar translational diffusion coefficient. For surface molecules, *N*_*hit*_ is adjusted accordingly based on the boundary edge length *L*. The total boundary crossing of molecules can be described as a Poisson process. As such, the number of molecules that cross the boundary each time step is drawn from a Poisson distribution with a rate *N*_*hit*_.

If a molecule enters the simulation box close to another boundary, the distance traveled parallel to the plane must be considered to resolve collisions with the mesh, although currently, PyRID does not account for this. For small integration time steps and meshes that are further than 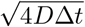 away from the simulation box boundary, the error introduced should, however, be negligible.

Once the number of molecules and their distance away from the boundary are determined, they are distributed in the simulation box (see 9 for more details). Diffusion along each dimension is independent, allowing for a random point to be selected uniformly distributed on the respective plane. For triangulated mesh surfaces, triangles are picked randomly, weighted by their area. Sampling uniformly distributed random points in a triangle is done using the method presented in [35].

Note that interactions between virtual molecules can generally not be taken into account by such methods such that fixed concentration boundary conditions only result in the same inside and outside concentrations if no molecular interactions are simulated. In addition, molecules that are close to a boundary will neither interact nor react across that boundary with any virtual molecule.

### 2.5 Berendsen barostat

In certain cases, it may be necessary to perform simulations in the NPT ensemble (constant number of particles, pressure and temperature), for instance, to release the system from stresses, compute interfacial properties of fluids [36], or compute phase diagrams using direct coexistence methods [13, 37]. The Berendsen barostat is a commonly used method for implementing NPT simulations, as it is simple to implement and results in the correct target density of the system. However, it does not sample from the correct statistical ensemble distribution; therefore, fluctuations in pressure tend to be too small. The Berendsen barostat scales inter-particle distances to adapt the forces and consequently altering the system pressure. PyRID supports simulations in the NPT ensemble using the Berendsen barostat.

### 2.6 Reactions

This section introduces chemical reactions of molecules supported by PyRID (Figure 3). These reactions can include for instance post-translational modifications, ligand binding, or ATP interactions. In general, the reactions can be defined on the molecule or bead/particle level, and are categorized as uni-molecular (first order), bi-molecular (second order), or zero order reactions. Uni- and bi-molecular reactions can involve multiple reaction paths, each belonging to a different reaction type.

**Figure 3:**
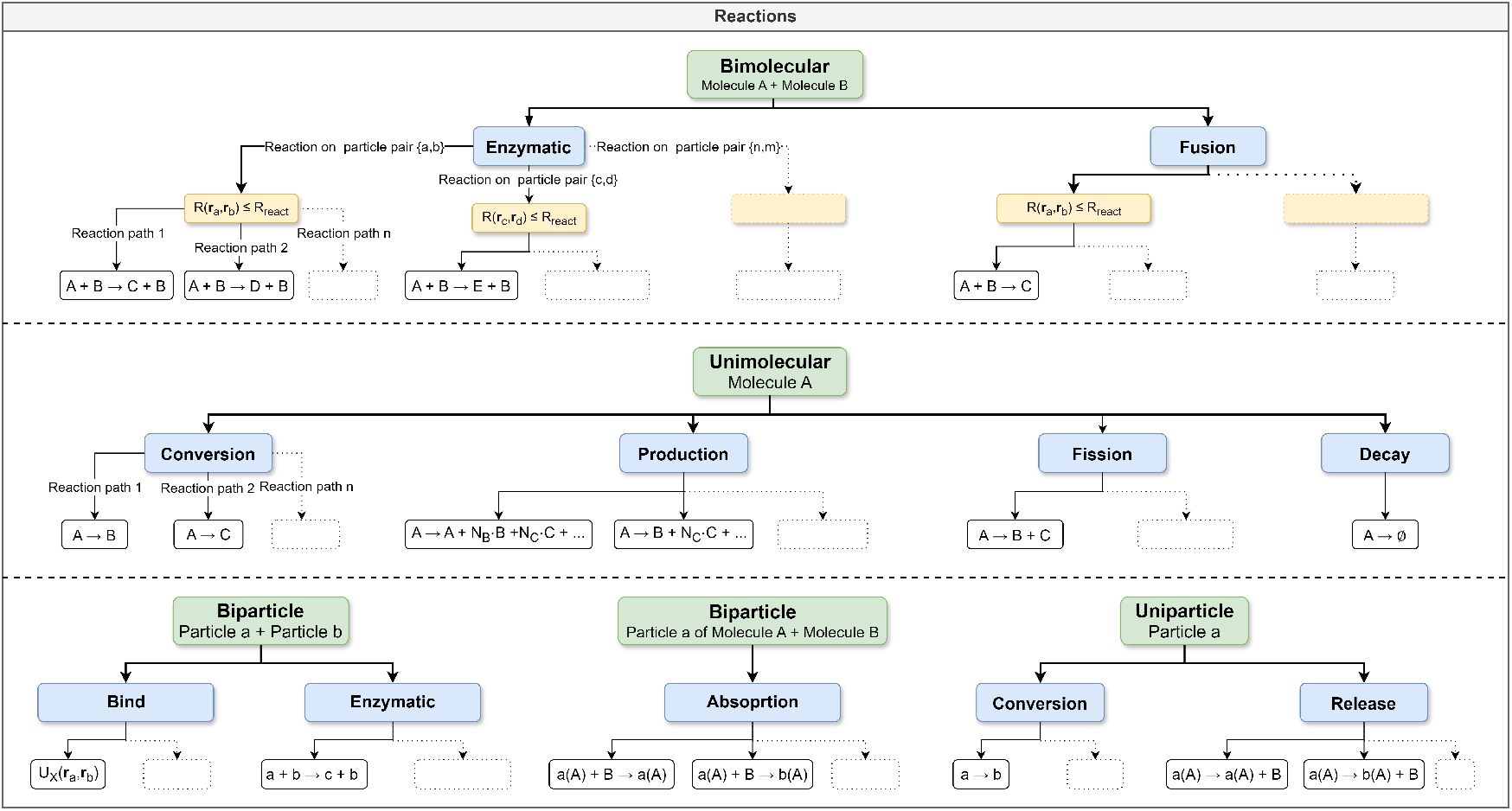
This tree graph provides an overview of the supported bi- and unimolecular reactions. A distinction is made between reactions that affect the molecules (upper case) and those affecting the particles (lower case). Both bimolecular and biparticular reactions are analyzed on the basis of particle pair distances *R*(*r*_*i*_, *r*_*j*_), since only these are calculated during the simulation and not, for example, the distances between the molecule’s geometric centers. Multiple reactions can be defined for the same particle pair, particle or molecule (e.g. a fusion and an enzymatic reaction), and multiple reaction paths can be defined for each of these reaction types, leading to various possible reaction outcomes or products.

#### Unimolecular reactions

Unimolecular reactions, such as decay, fission, and conversion, can be efficiently simulated using a variation of the Gillespie Stochastic Simulation Algorithm (SSA) [38, 39, 40]. This method samples the time point of the next reaction from the expected molecule lifetimes probability distribution, assuming no interfering events such as a bi-molecular reaction occur in between. The Gillespie SSA is beneficial, as it is exact and evaluates a reaction only once, instead of each time step. For a single molecule having *n* possible reaction paths each with a reaction rate 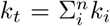 be the total reaction rate. Let *ρ*(*τ*)*dτ* be the probability that the next reaction occurs within [*t, t* + *τ* + *dτ*), which can be split into *g*(*τ*), the probability that no reaction occurs within [*t, t* + *τ*) and probability that a reaction occurs within the time interval *dτ*, which is given by *k*_*t*_*dτ*. Thereby,

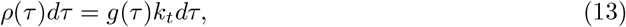

where 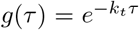[38]. From the above equation we find 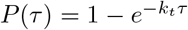by integration. To sample from this distribution, we can use the inverse distribution function:

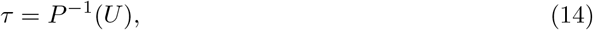

where *U* is uniformly distributed in (0, 1). With 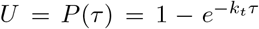, we find 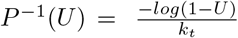 Thereby, we can draw the time point of the next reaction from:

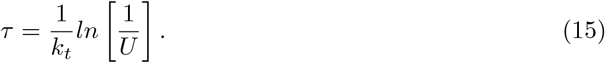

With the above method, we accurately sample from the distribution of expected molecule life-times 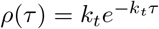. These reactions can, amongst others, describe protein degradation, ligand unbinding, disassembly of protein complexes, and changes in folded protein states. In PyRID, uni-molecular reactions can occur on either the molecule or particle level, depending on the definition of the reaction. If a conversion or decay reaction is defined on a molecule, the complete molecule is exchanged with the product or removed in the case of a decay reaction. However, if defined on a particle, only the particle is affected. PyRID offers three types of fission reactions termed - fission and production reactions defined on molecules, and release reactions defined on particles/beads (Figure 4).

**Figure 4:**
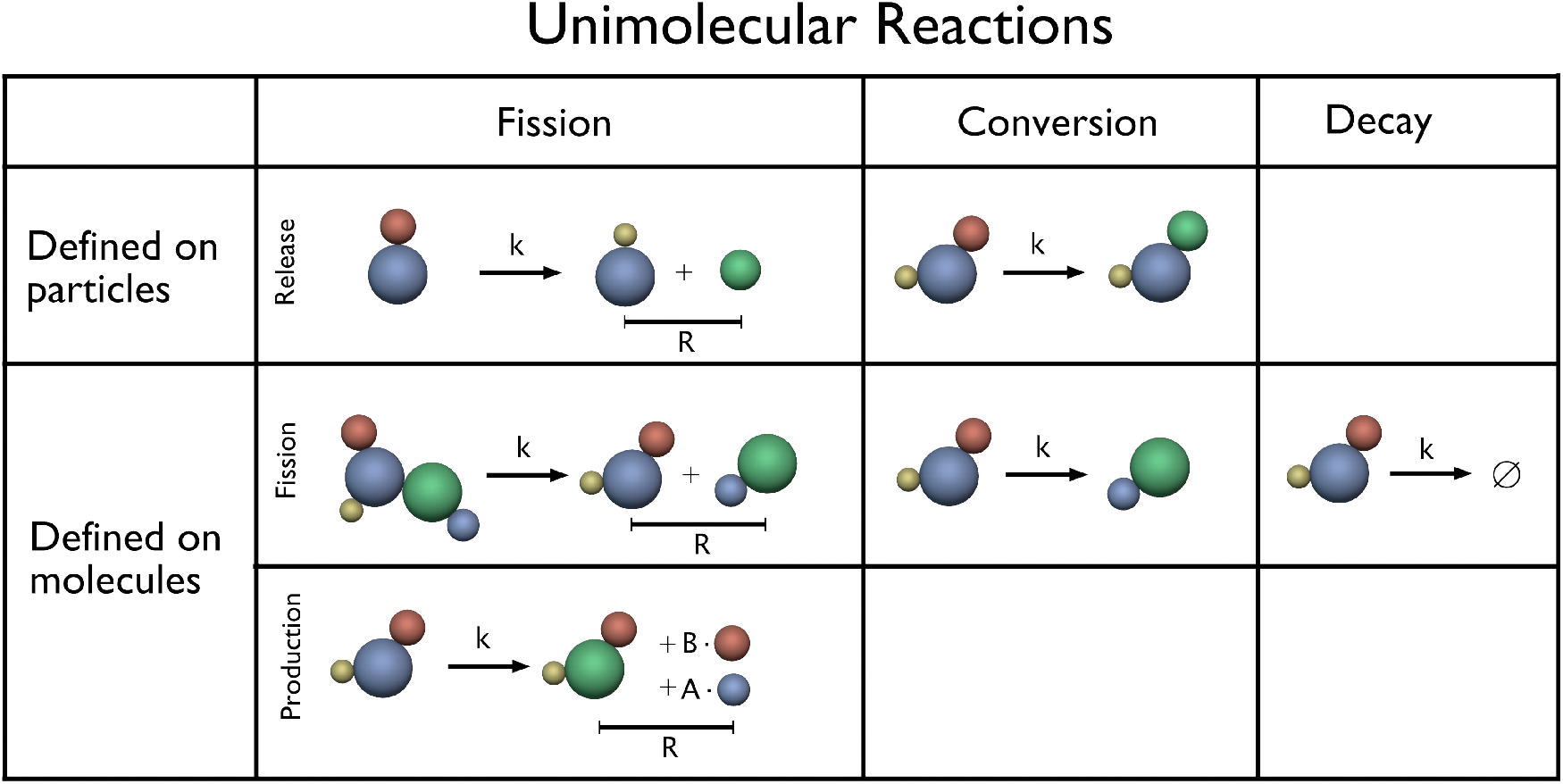
Schematic of unimolecular reactions. Unimolecular reactions can be either defined on a particle or on a molecule type. Release reactions are a sub-category of the fission reaction. Here, a molecule (rigid bead model) is released from another molecular particle/bead. Release reactions can be used, for example, to model the release of a ligand from a specific molecule’s binding site. The production reaction enables a fission reaction with more than two products. It can be used, for example, to model the influx of ions into a compartment via an ion channel.

#### Bi-molecular reactions

Due to the uncertainty of molecule encounters, bi-molecular reactions require a different approach than uni-molecular reactions. PyRID utilizes the Doi reaction scheme [41]. In this scheme, two molecules can only react if the inter-molecular distance |*r*_*ij*_| is below a reaction radius *R*_*react*_. The probability of having at least one reaction is then given by

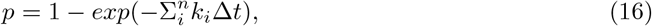

where *n* is the number of reaction paths.

PyRID supports various bi-molecular reactions, including fusion reactions, enzymatic reactions, and binding reactions, which can be defined on molecules and/or particles (Figure 5). For more details see the PyRID documentation.

**Figure 5:**
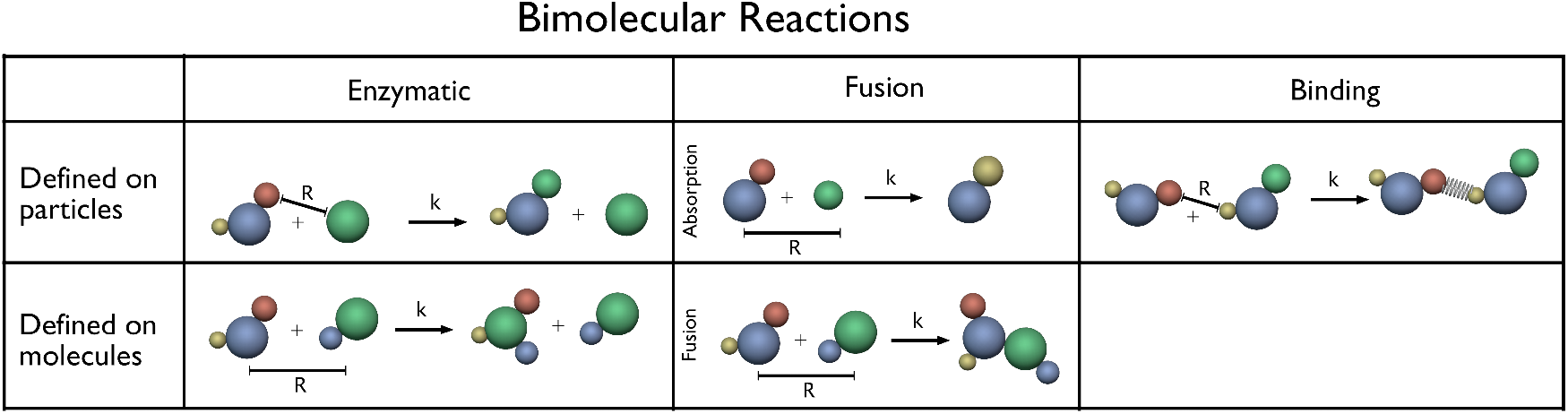
Schematic of bimolecular reactions. Bimolecular reactions can be either defined on a particle or on a molecule type. The absorption reaction is a sub-category of the fusion reaction. It has been introduced as the inverse to the release reaction (Figure 4) and can be used to model, e.g., ligand binding. Here, a molecule is absorbed by the bead/particle of another molecule and thus removed from the simulation. Binding reactions introduce an energy potential between two particles/beads. Short-range interactions can be used, for example, to model patchy particles (Figure 15).

Please note that in PyRID surface reactions between molecules on a surface are based on their Euclidean distance, which may not accurately reflect the local surface curvature. Accurately calculating geodesic distances on mesh surfaces is computationally expensive, and fast approximations such as Dijkstra’s algorithm may not provide good approximations for points close to the source. In addition, as reaction radii are typically small compared to the mesh size and surface curvature, deviations in calculations are expected to be small in most cases. Nonetheless, recent progress has been made in this area (e.g., 42, 43, 44) and could be implemented into PyRID in future versions.

### 2.7 Distribution of molecules

#### Volume molecules

When dealing with mesh compartments and accounting for molecule excluded volume, distributing molecules in the simulation volume becomes a challenging task. A standard approach involves loosely distributing molecules and shrinking the volume until reaching a target density. This approach could be transferred to a system with mesh compartments.

However, here, we might also care about the exact dimensions of the compartment so that the synchronous convergence of the desired dimensions and the molecular density becomes an additional challenge. Alternatively, the Metropolis Monte Carlo method can be used [45], but it is time-consuming. Instead, PyRID employs the Poisson-Disc sampling algorithm [46], which is computationally efficient and straightforward to implement but has a density limit of 30% and is only suitable for spherical molecules. For highly aspherical molecules, Monte-Carlo sampling is used, and overlaps are resolved via a soft repulsive potential. The Berendsen barostat at high pressure can also drive the system to a high-density state if no mesh compartments are used.

The Poisson-Disc sampling algorithm consists of initializing a grid, creating a sample point, and inserting it into a list of active elements. Random points around the annulus of the active sample points are created, and new sample points are accepted and inserted into the grid and active list if no other sample points exist within the radius. If no new sample point is found after *k* trials, the active sample point is removed from the active list. PyRID extends this algorithm to account for polydisperse particle distributions (see Figure 6 A).

**Figure 6:**
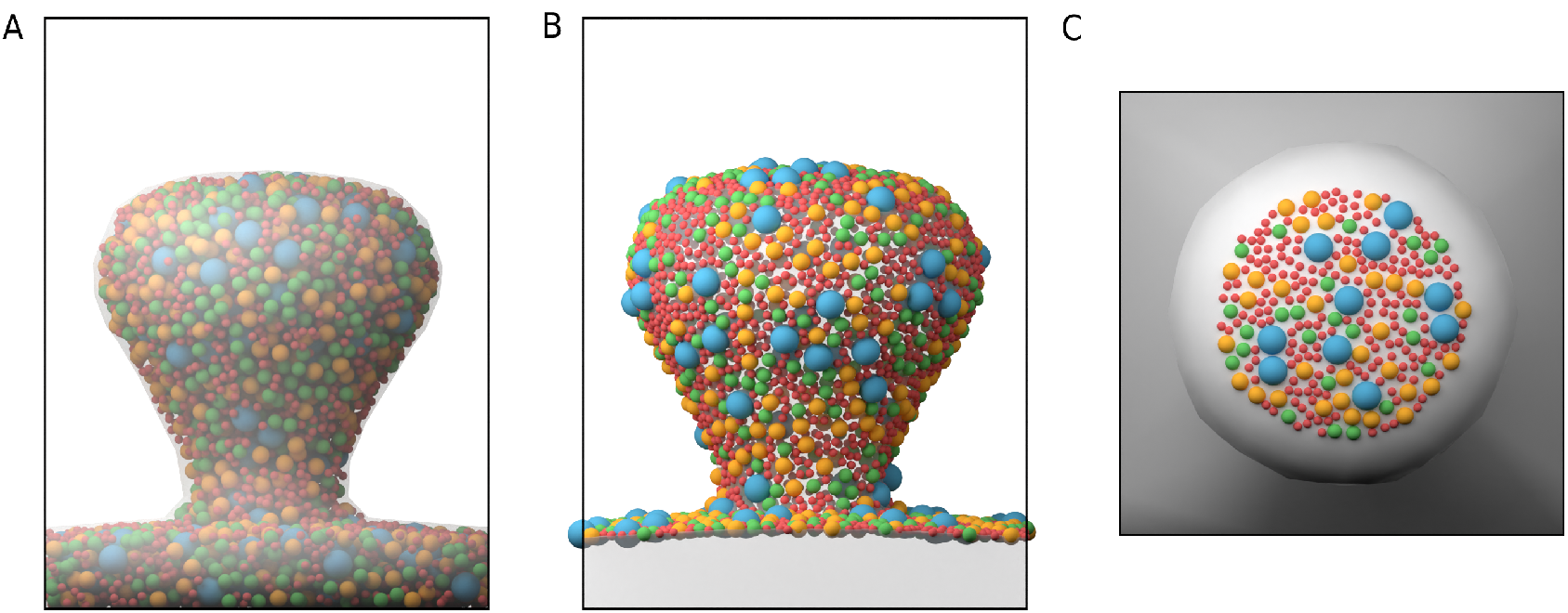
Poisson Disc Sampling of polydisperse spheres. (A) Example distribution for three different sized particle types confined to the volume of a mesh compartment. (B) Poisson Disc sampling for surface molecules. (C) Poisson Disc sampling for surface molecules being restricted to a surface region that is defined by a triangle face group.

#### Surface molecules

We use an algorithm introduced by [47] to distribute molecules on the surface of a mesh compartment. The algorithm involves generating a sample pool, dividing space into cells, randomly selecting cells and samples, checking distances between occupied and unoccupied samples, and updating the number of samples for each cell. The process is repeated until the desired number of molecules is reached or a maximum number of trials is reached. PyRID also allows the user to assign surface regions for molecule distribution. Example is shown in Figure 6 B & C.

### 2.8 Fast algorithms for Brownian dynamics of reacting and interacting particles

The simulation performance in PyRID is optimized using Numba’s jit compilation. Additionally, PyRID employs a data-oriented design and dynamic array data structures to efficiently track molecules and their reactions. PyRID requires a data structure capable of efficiently handling the constant changes in the number of particles and molecules within the system due to reactions and events. Similarly, the molecular reactions occurring at each time step must be listed and evaluated efficiently. To achieve this, PyRID utilizes two variants of dynamic array data structures, namely the tightly packed dynamic array and the dynamic array with holes, both well-suited for these tasks.

#### The tightly packed dynamic array (dense array)

A tightly packed dynamic array is a variant of dynamic arrays used in PyRID that enables efficient deletion of elements via a pop and swap mechanism. Unlike standard numpy arrays, lists, or C++ vectors, the tightly packed dynamic array does not create a new array every time an element is deleted or appended, which significantly reduces computational cost. This is achieved by increasing the array size by a multiplicative factor and keeping track of the length and capacity of the array. However, this method requires keeping track of the location of elements in the array as they move around during deletion or swapping, which is accomplished by maintaining a second array (see Figure 7 left).

**Figure 7:**
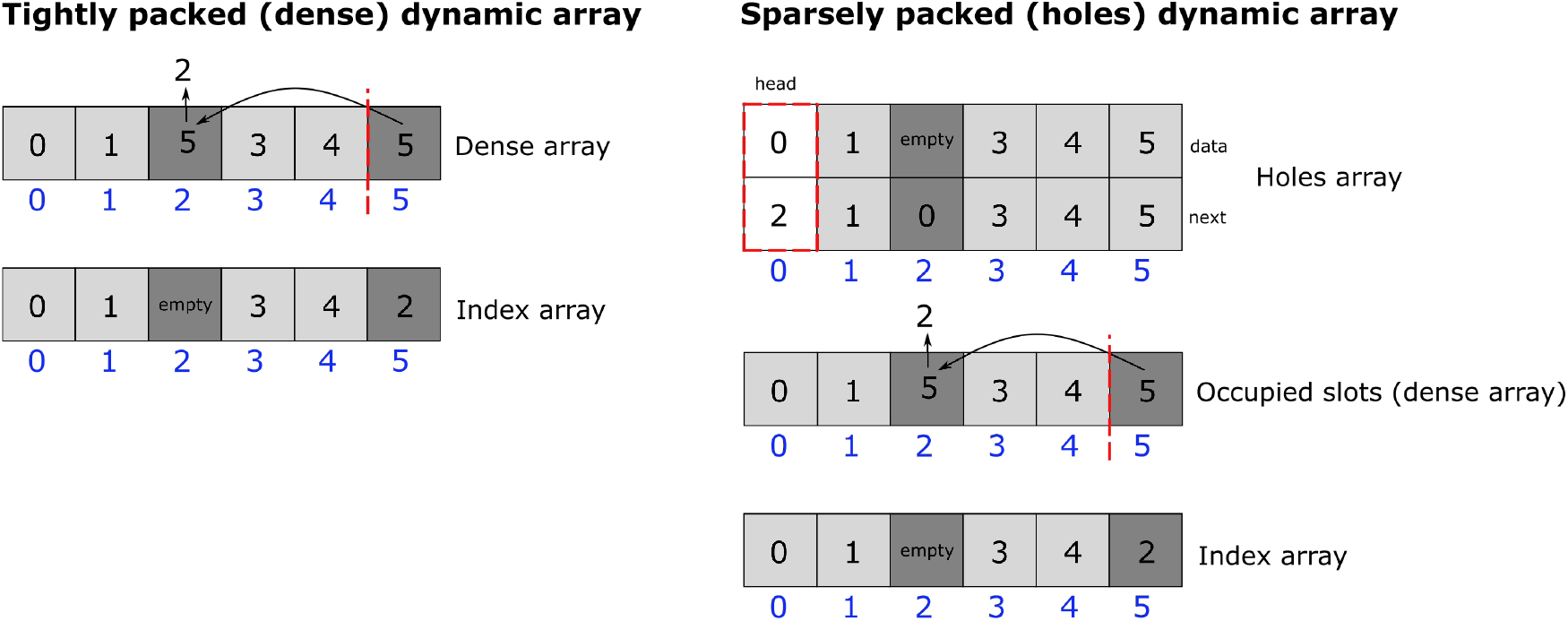
Dynamic arrays. (left) Tightly packed dynamic array. (right) Sparsely packed dynamic arrays.

#### The dynamic array with holes

The dynamic array with holes method (Figure 7 right) is used to store molecules and particles, where holes are created to enable efficient deletion, tracked via a free linked list. This approach preserves the original indices and allows for fast element access. However, if the array is sparse, it may not be cache-friendly and iterating over elements becomes complex. To address this, a tightly packed array is added to store occupied slot indices for efficient iteration. The strength of this method lies in its ability to preserve original indices while enabling fast element access and efficient deletion.

#### Dynamic arrays used for reaction handling

The data structure required to organize reactions is more complex than a simple dynamic array as used for rigid body molecules and particles. To efficiently add, delete single and all reactions of a particle, and return a random reaction from the list, a combination of dynamic arrays and a hash table is used. A doubly linked list is embedded into a dynamic array with holes to connect all reactions of a particle, and its head is saved in a hash table. A 4-dimensional next and previous pointer is required to keep track of the educts in a reaction. A separate dynamic array is used to keep track of all remaining reactions in a tightly packed format for easy random selection. The proposed data structure avoids introducing biases and efficiently performs the required operations.

### 2.9 Polydispersity

In molecular dynamics simulations, the polydispersity of particle radii can pose a problem, particularly when using minimal coarse-graining approaches with low granularity. This can lead to classical linked cell list algorithms becoming inefficient. The most computationally expensive task in such simulations is typically the calculation of pairwise interaction forces, which requires determining the distance between particles. Naively iterating over all particle pairs results in a quadratic increase in computation time with the number of particles in the system. The linked cell list approach provides a straightforward solution to this problem [45]. By dividing the simulation box into cells of appropriate size, the computational complexity can be reduced to a more linear increase with N. The number of cells must be chosen such that the side length of the cells in each dimension is larger than the maximum cutoff radius for pairwise molecular interactions. The resulting computation time is of the order of *N* instead of quadratic *N* ^2^, assuming a mostly homogeneous distribution of molecules.

However, this approach does not efficiently handle polydisperse particle size distributions, which can result in inefficient calculations when using minimal coarse-graining approaches with low granularity. To address this issue, a hierarchical grid approach has been introduced by [2]. In this approach, which is used in PyRID, each particle is assigned to a cell grid based on its cutoff radius, resulting in different levels or hierarchies, each with a different cell size. Although this approach requires more memory, it significantly reduces the number of distance calculations required for larger particles and takes advantage of Newton’s third law. The algorithm involves assigning particles to cells on their respective level, performing distance checks for nearest neighbor cells on the same level, and then doing a cross-level search using distance checks only on lower hierarchy levels. However, this approach may require iterating over many empty cells during the cross-level search.

### 2.10 Simulation algorithm

The simulation loop of PyRID begins by updating the position of all particles, followed by determining the particle pair distances and calculating the forces, which are then added to the reactions list. Reactions are performed next, allowing new particles to enter the system through fusion or fission reactions. To save computation time, the inter-particle distances and forces are not updated, except for binding reactions where the force is calculated at the time of reaction execution. In the next iteration, the reaction products diffuse freely without experiencing any forces yet, and the trajectory of particles that were already in the system, remains unchanged due to the presence of new molecules. The placement of product molecules does not resolve any collisions with nearby molecules, introducing a small error. Fission reactions have a similar error regardless of whether forces are updated or not. For fusion reactions, the error is slightly different as the educts leave behind an empty space that will be filled with new molecules. To avoid issues with a dense system, the integration time step should be decreased.

An alternative approach would be to calculate all forces and reactions at the beginning of each iteration before updating molecule positions. However, this method can introduce bias, especially for bimolecular reactions. PyRID’s approach avoids bias when interactions are not considered. ReaDDy updates the neighboring list for molecules twice per iteration, which could double computation time if not optimized. PyRID evaluates forces and reactions in one loop over all particle pairs in the neighboring list, with pair distances calculated only once. Future work could optimize the update of pair distances after reactions are executed.

### 2.11 Visualization

For visualization, we have developed a blender addon for PyRID (Figure 8). In addition, PyRID can visualize the different reactions using graphs (Figure 9). Graph visualization in PyRID is build upon the pyvis library.

**Figure 8:**
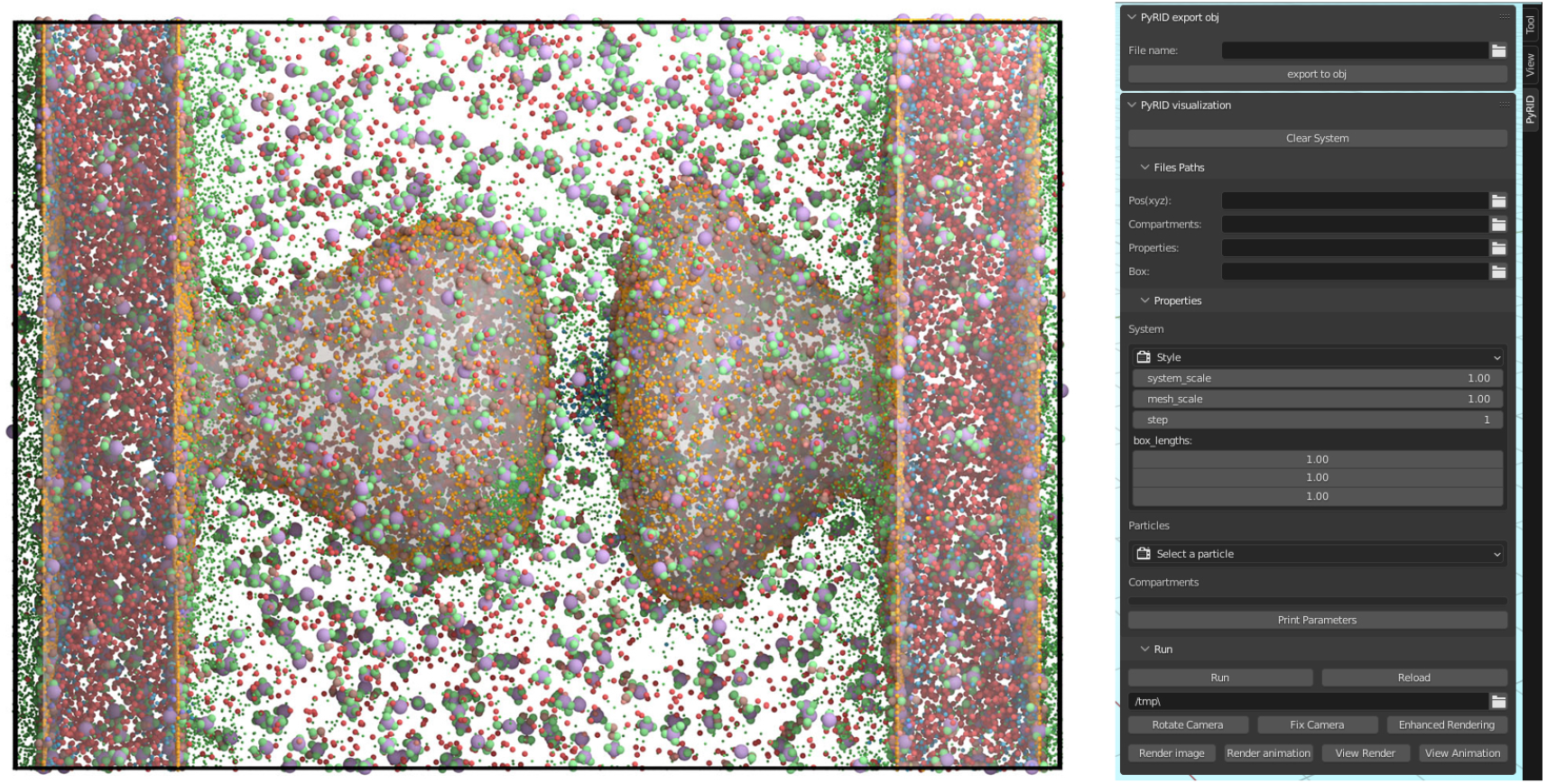
Visualization of molecule trajectories with PyRIDs Blender addon. Left: Example visualization with 50.000 particles. Right: GUI of the Blender addon.

**Figure 9:**
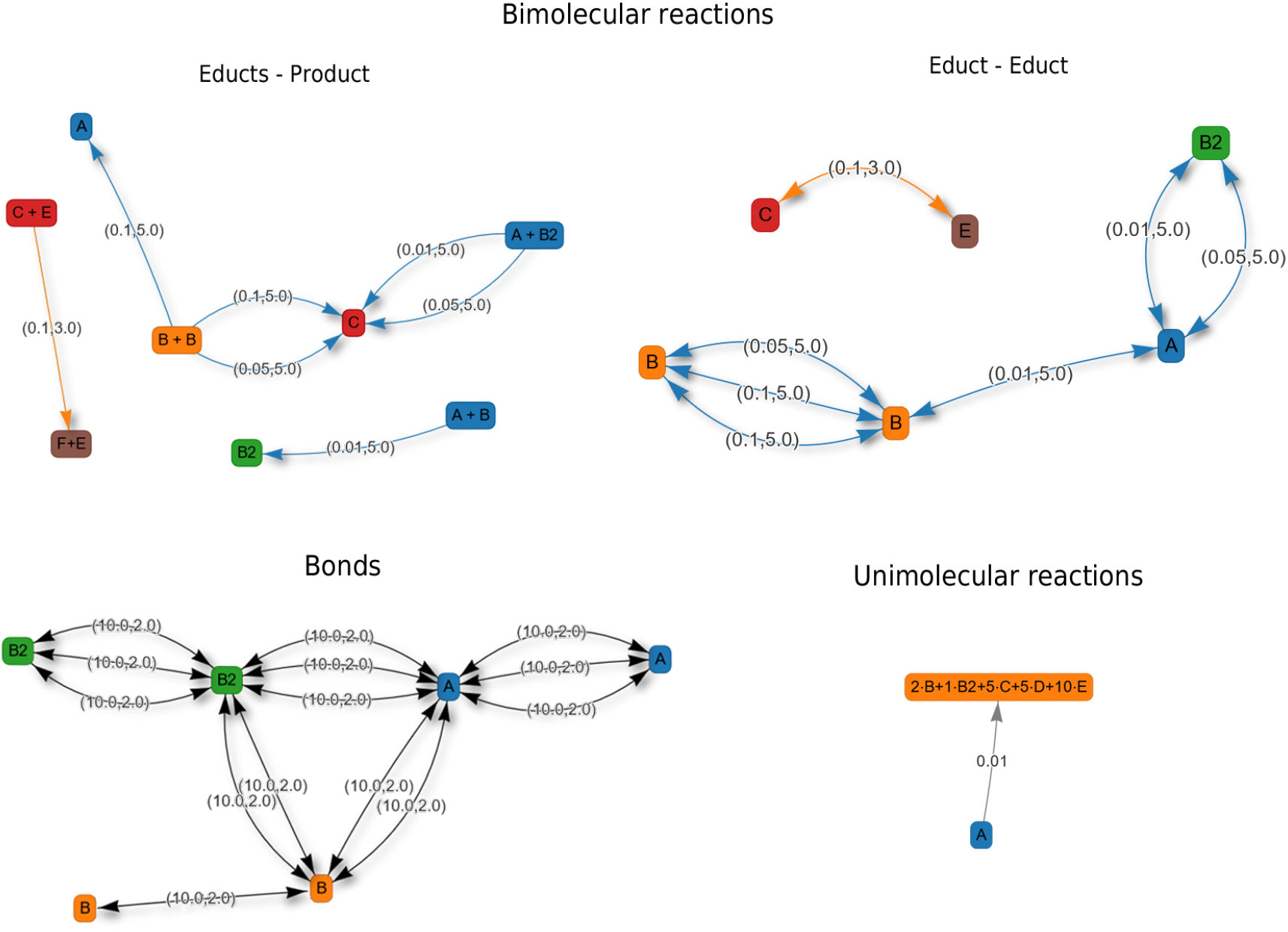
Examples of different reaction graphs. The “Educts - Product” graph shows the reactants and their respective products as well as the various possible reaction paths. The reaction rate and the reaction radius are shown in round brackets. The “Educt - Educt” graph only depicts the relationship between the different educts. The “Bonds” graph shows the binding reactions between particle pairs.

## 3 Results and Validation

### 3.1 Anisotropic diffusion

We verified the accuracy of our translational and rotational diffusion algorithms using the example from [22], where the diffusion tensors are not molecule-specific. We compared the mean squared displacement (MSD) and rotational time correlation with theoretical predictions, using a multi-exponential decay function for the latter [48]. The MSD is given by:

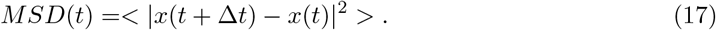

The rotational time correlation function is given by:

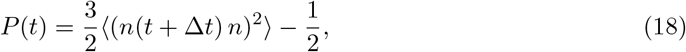

where *n*(*t*) is some unitary vector that describes the current orientation of the molecule at time point t. Figure 10 shows the comparison between simulation results and theoretical prediction.

**Figure 10:**
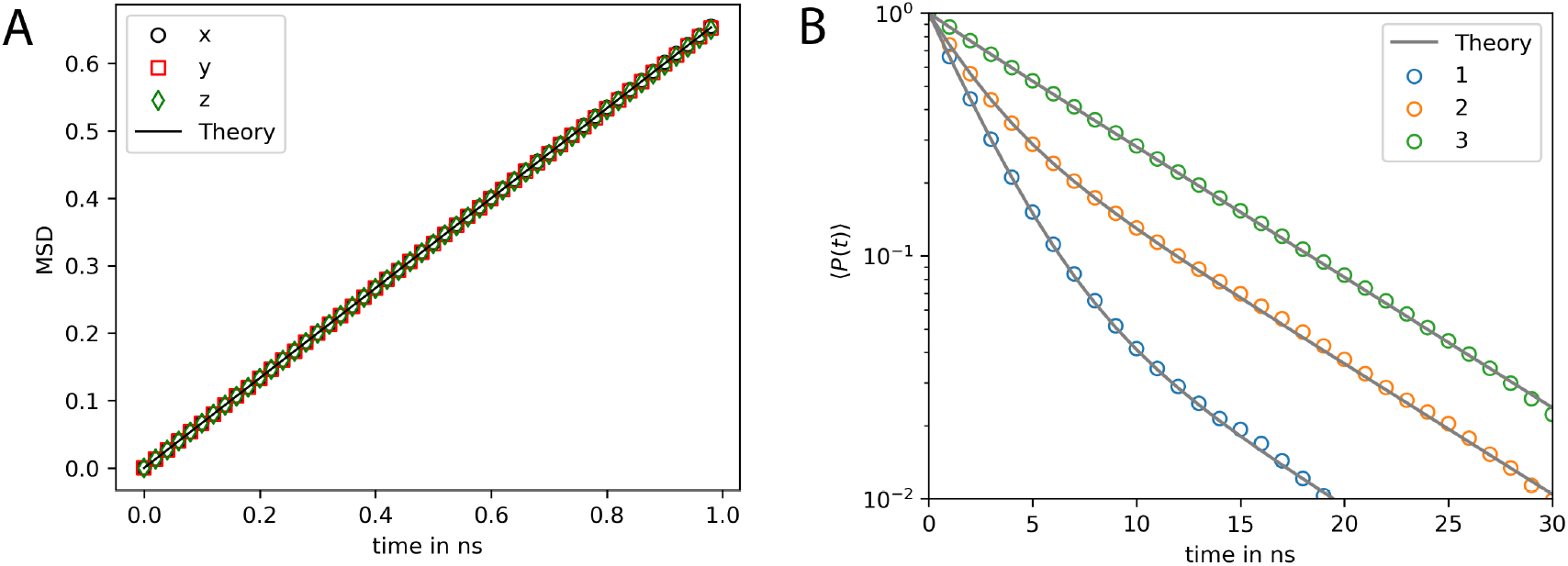
MSD and rotational relaxation times of a rigid bead molecule matches the theoretical prediction. (A) Mean squared displacement (MSD) of the rigid bead molecule computed with PyRID. The displacement in each dimension (colored markers) is in very good agreement with the theory (black line). (B) The rotational relaxation of the rigid bead molecule is also in close agreement with theory (gray lines) for each of the the rotation axes (colored markers).

### 3.2 Diffusion tensor of the igG3-protein

The methods described to compute diffusion tensors were previously only available in Hydro++, but we have now incorporated them directly into PyRID for efficient setup of rigid bead molecule systems. We tested our implementation using the igG3 protein model (Figure 11) provided in the Hydro++ documentation and obtained results that are in good agreement with Hydro++ at up to 4 decimal places.

**Figure 11:**
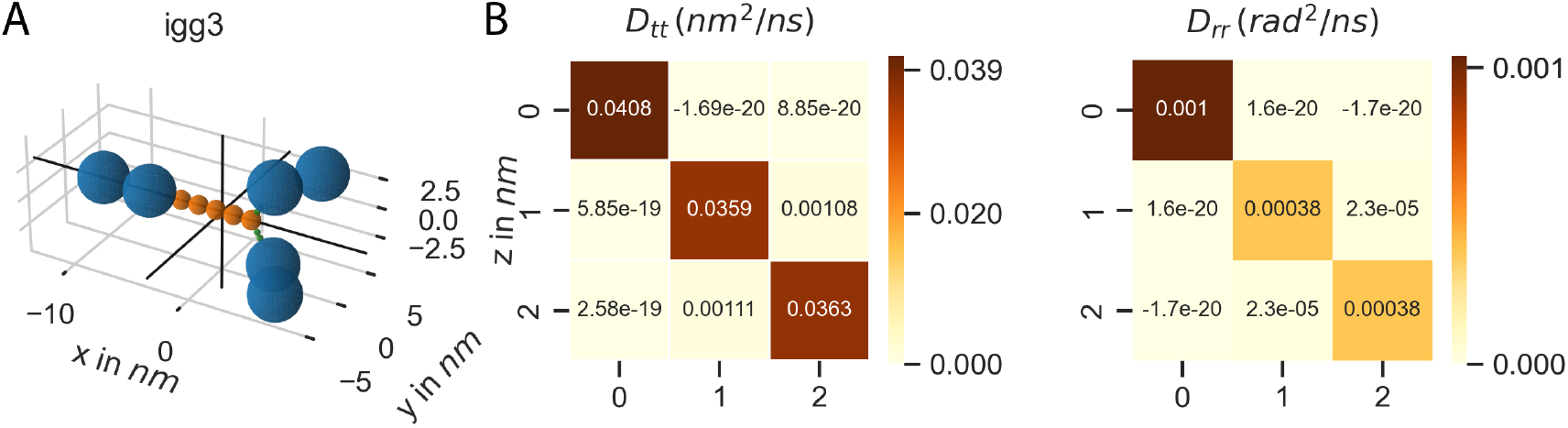
The diffusion tensor of igG3 calculated with PyRID. (A) Rigid bead molecule representation of igG3 as found in de la Torre and Ortega (2013). The origin of the coordinate system is set to the center of diffusion of the igG3 rigid bead model (bold black lines). (B) Translational and rotational diffusion tensor of igG3.

### 3.3 Choosing the right reaction rate and radius

In molecular simulations, accurately choosing reaction parameters such as reaction rate (*k*) and reaction radius (*R*_*react*_) is crucial for capturing realistic system behavior. Hereby, it is important to choose *R*_*react*_ carefully to avoid reactions with non-nearest neighbors in dense systems.

However, *R*_*react*_ should also not be smaller than the average change in the distance between molecules, which is given by 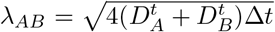 where 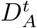 and 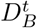 are the translational diffusion constants of two molecular species A and B. Otherwise, a molecule might pass many reaction partners in between two time steps where the bi-molecular reactions are not evaluated [49]. Alternatively, as proposed by [50], one can use the electrostatic interaction length scale to define *R*_*react*_.

A description of the reaction kinetics in terms of a system of differential equations assumes a well mixed system. Therefore, the simulation results are also only directly comparable with the ODE approach, if the reactions are reaction rate limited, not diffusion limited such that the system has enough time to equilibrate in between reactions. In a simple *A*+*B→C* reaction with diffusion-limited kinetics, the local concentration of educts only gradualy decreases as mixing with products by diffusion is slow in comparison to the microscopic reaction rate. As a consequence, the probability of encounter for the remaining educts also only decreases slowly. Educts and products are heterogeneously distributed, the system is not well-stirred. In contrast, in a well-stirred system, the concentration of educts globally and locally decreases, reducing the probability of educt encounters. Since the assumption of a well-stirred system is no longer valid for diffusion-limited kinetics, the reaction kinetics in the ODE approach are slowed down compared to the stochastic particle based simulation (Figure 12).

**Figure 12:**
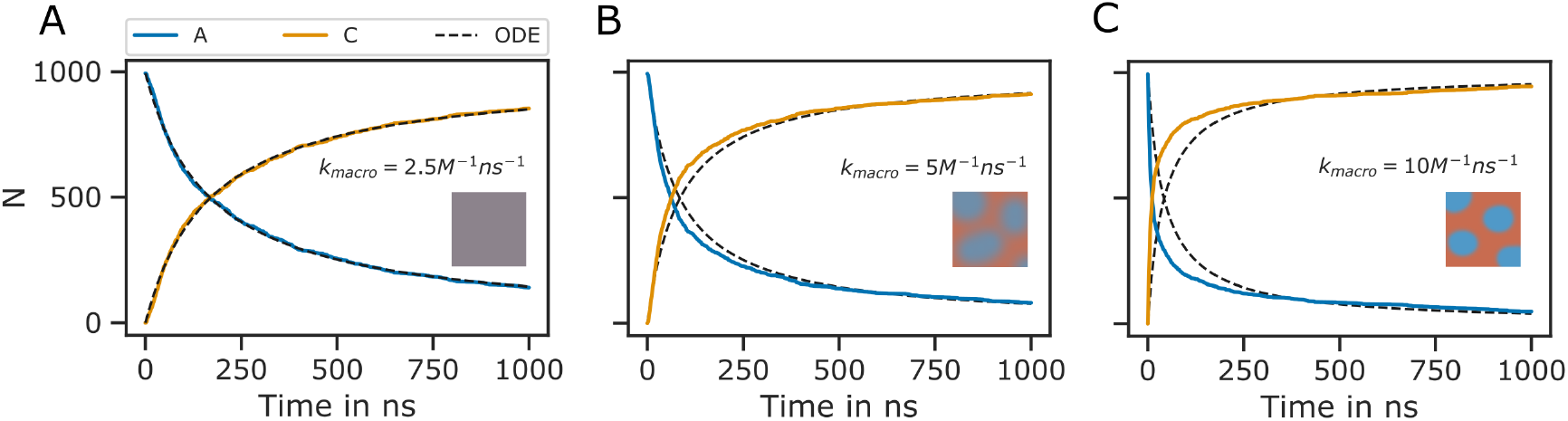
Diffusion limited bi-molecular reactions are not accurately described by ODEs. Shown is the minimal system 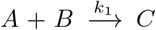 with *R*_*react*_ = 4.5*nm* and *σ*_*A*_ = 3*nm, σ*_*B*_ = 4.5*nm, σ*_*C*_ = 3.12*nm*. The same system has been used for validation of ReaDDy [11]. The ODE approach to the description of the reaction kinetics assumes a well mixed system. If the reaction rate is small, the system has enough time to equilibrate in between reactions and the ODE approach (black dotted lines) and the particle-based SSA approach (colored lines) match. (A). However, with increasing reaction rates (B-C), the system becomes heterogeneous over time, resulting in regions of varying educt concentrations. Initially, stochastic simulations exhibit faster kinetics than ODE predictions (B, C). However, when a critical mass of educts have reacted,slow diffusion causes kinetics to deviate from ODE predictions, necessitating two exponential functions instead of one. The slow down effect is especially prominent in B, C at around 500 ns. The reaction kinetics are therefore better described by two exponential functions instead of one.

The reaction rate determining the probability that a reaction occurs within a given time step in particle based simulations is not the same as the reaction rate in the ODE approach. In line with [50] we refer to the latter as the macroscopic rate and other as the microscopic rate. To compare the particle based simulations to the ODE approach, the microscopic reaction rates need to be converted to the corresponding macroscopic rates or vise versa, which can be done by taking into account the reaction radius and the diffusion constants of the reacting species.

Given a reaction radius *R*_*react*_, we would like to know at what reaction rate *k* a simulation would match an experimentally measured macroscopic reaction rate *k*_*macro*_. For two non-interacting molecule species A and B with translational diffusion constants 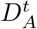and 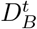 and *λ*_*AB*_ *<< R*_*react*_, *k*_*macro*_ is given by [49]:

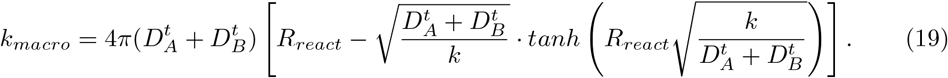

Eq. 19 can be solved numerically for *k*.

PyRID has a build in method to calculate the microscopic reaction rate *k* and the macroscopic reaction rate *k*_*macro*_ from Eq. 19.

### 3.4 Bi-molecular reactions between rigid bead molecules

The use of single particles to represent complex molecules neglects their structure and potential for bi-molecular reactions via different sites. PyRID addresses this limitation by enabling the simulation of reactions between complex molecules with different reaction sites represented by beads/patches in the rigid bead molecule’s topology. Bi-molecular reactions can be defined on particles or molecules, but in the latter require linkage to a particle type pair for distance computation. Successful reactions on molecule types occur if the two particles are within the reaction distance. As an example, we consider molecules *A* and *B* that are represented by two beads each *a*_1_, *a*_2_ and *b*_1_, *b*_2_, and reaction *A* + *B↔C* as well as *A* + *B→D*. We now may define reactions for different pair permutations of the available beads like the following:

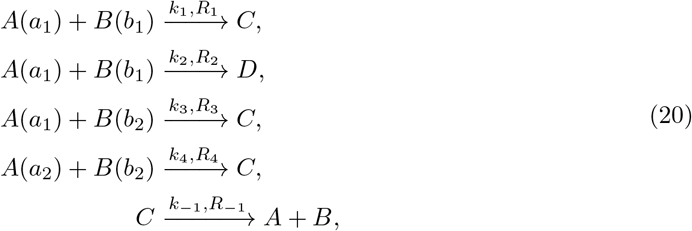

where *k*_*i*_ are the microscopic reaction rates and *R*_*i*_ the reaction radii. For better visualization, see Figure 13 A and B. Whereas for the particle pairs (*a*_1_, *b*_2_) and (*a*_2_, *b*_2_) only one reaction pathway is defined respectively, for the particle pair (*a*_1_, *b*_1_) a second reaction path has been defined for the fusion of molecules *A* and *B* to molecule *D*. We may also describe this system in terms of a system of ODEs:

**Figure 13:**
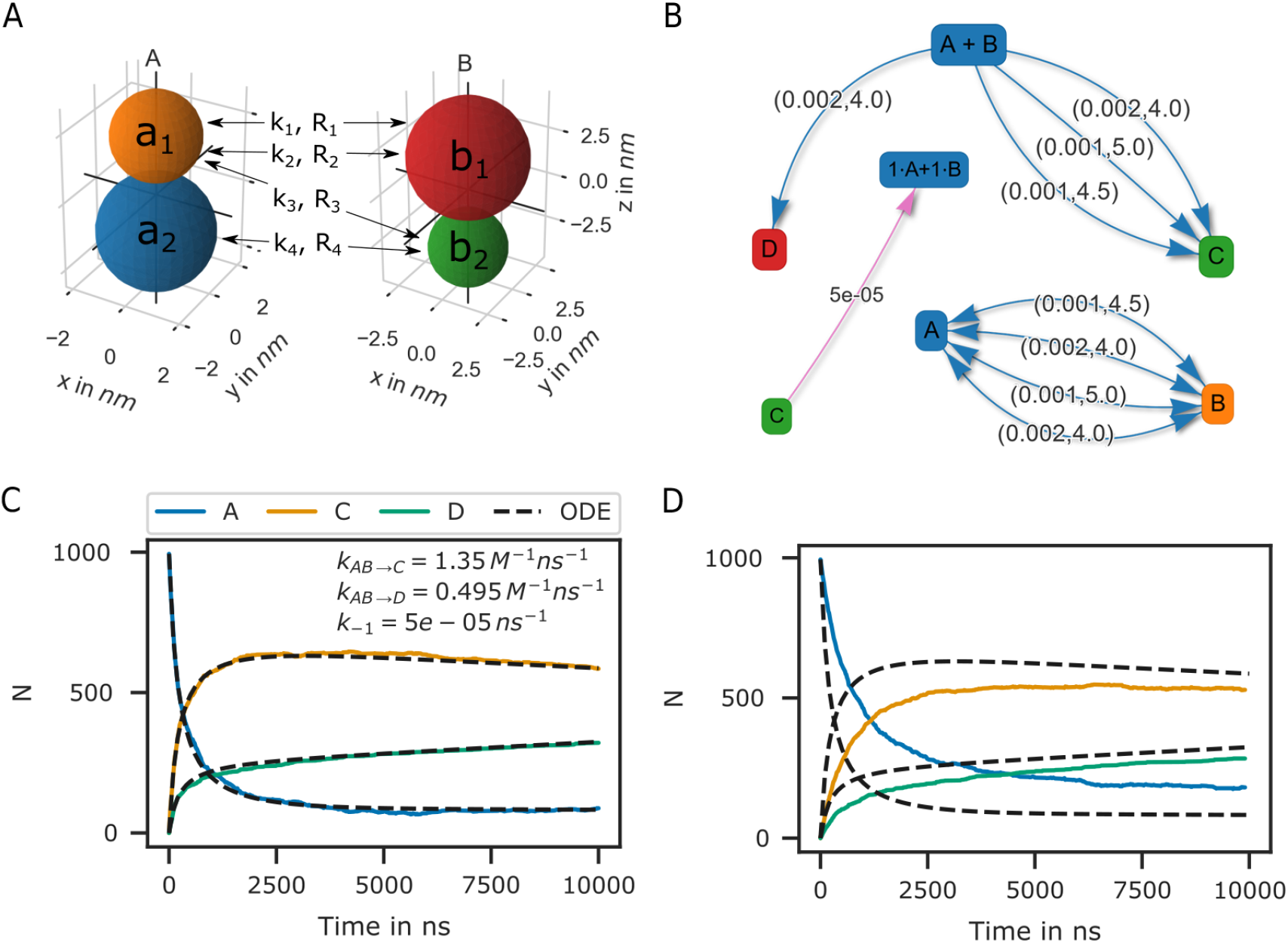
Bi-molecular reaction between two rigid bead molecules. (A) Depiction of the two rigid bead molecules and the different reactions defined on their respective particles/beads. (B) Reaction graphs showing the different reaction paths for the fusion reactions and as well as the fission reaction. The lower right graph depicts the different reaction paths between the two educts *A* and *B* without specifying the products. In total there are 4 paths. (C) If not accounting for any repulsive interaction between molecules *A* and *B*, the simulation results are in good agreement with the ODE description. (D) If we account for the excluded volume of the molecules by a repulsive interaction potential, the results of the particle simulation and the ODE no longer match.

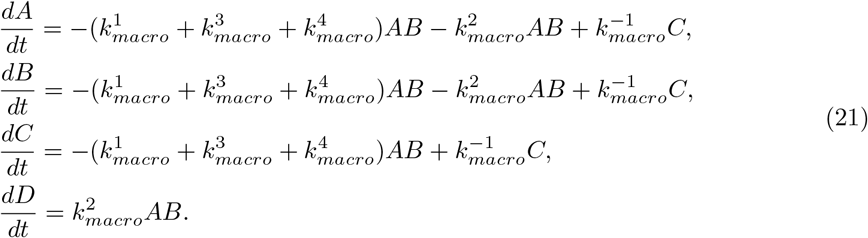

The macroscopic rate constants *k*_*macro*_ can be calculated from Eq. 19. For complex molecules, however, Eq. 19 is inadequate for calculating the macroscopic rate constants, as it only accounts for the translational diffusion constant of the molecule center and not the rotational motion. For simple molecules such as the bead in our example, the Brownian dynamics simulation results align closely with the ODE formulation (Figure 13 C), indicating a similar motion to that of a single spherical particle. However, for more complex molecules with anisotropic reaction volumes, a deviation from this approximation is expected. Although the rigid bead model may not seem particularly advantageous over other stochastic simulation algorithms for the above example, it already becomes useful when considering excluded volume effects. By introducing repulsive interactions between the beads, we observe differences in reaction kinetics when compared to the ODE solution (Figure 13 D).

### 3.5 Hard sphere fluid

For validation purposes, a hard sphere fluid is advantageous due to the availability of analytical expressions for the radial distribution function and pressure.

#### Radial distribution function

In Figure 14 B, the radial distribution function is shown for a hard sphere fluid simulated with a harmonic repulsive interaction potential and a sphere diameter of 1 nm. The simulation result agrees well with an analytical expression for the hard sphere radial distribution function from [51].

**Figure 14:**
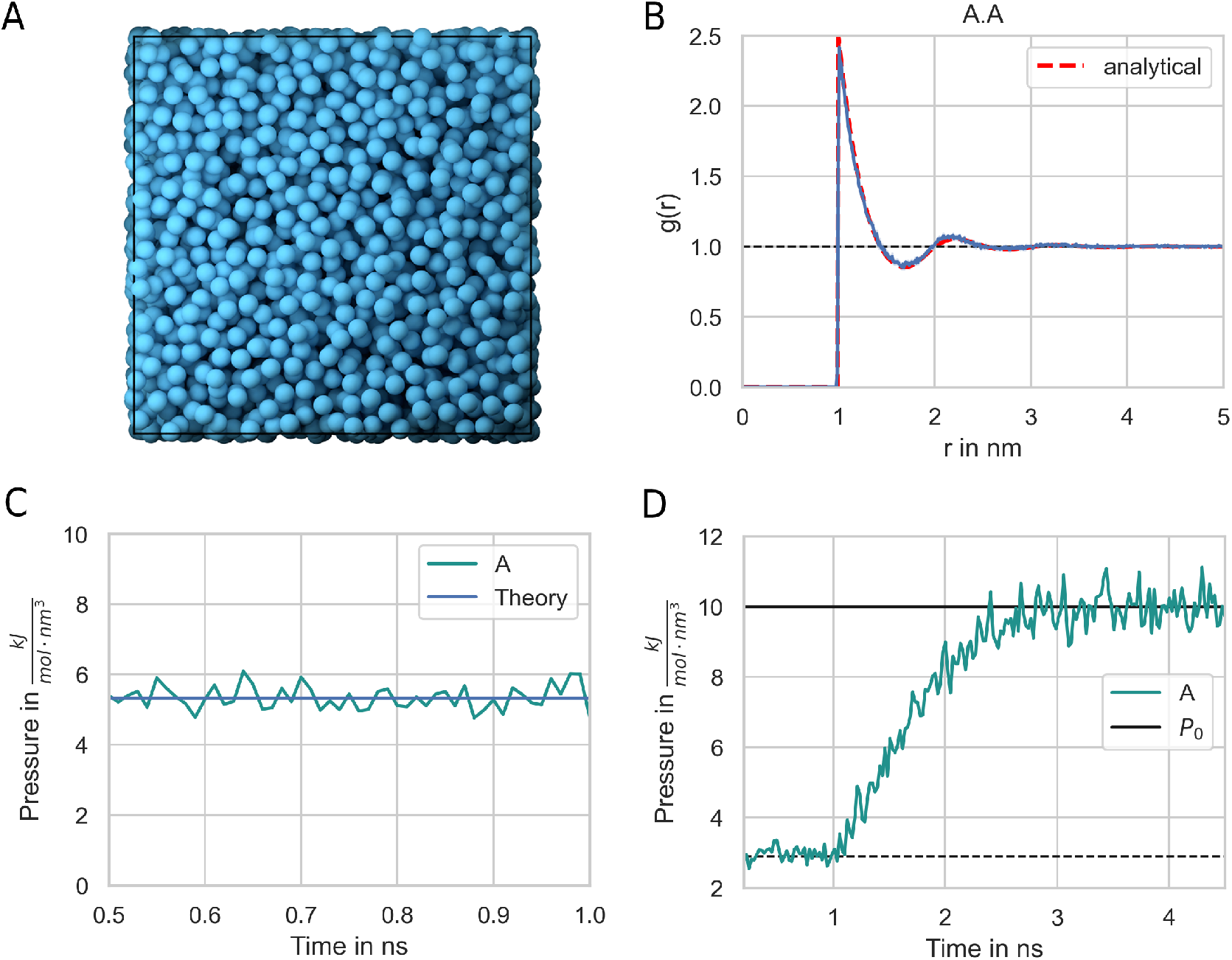
Hard-sphere radial distribution function. (A) The system is set up with a packing fraction of *η* = 0.3. The particle diameter is set to 1 nm and pair interactions occur via a harmonic repulsive potential. (B) The resulting radial distribution function (blue line) is in close agreement with theoretical prediction (red line). (C) The pressure of the hard-sphere fluid obtained from simulations is also in close agreement with theory (blue line, 51). (D) A hard-sphere fluid NPT ensemble simulation. From time 0.5 ns, the Berendsen barostat is activated and drives the system to the target pressure. *P*_0_ = 10 *kJ/*(*mol nm*^3^) = 16.6 *MPa* = 166 *bar*

#### Pressure

A hard sphere fluid can serve as a reliable means of validating pressure calculations. The pressure can be expressed analytically in terms of the radial distribution function at contact and the second virial coefficient [52], with the Percus-Yevick equation offering an approximation for the radial distribution function at contact [53]. The simulation results for the pressure of a hard sphere fluid closely match theoretical expectations (Figure 14 C), and the system reaches the target pressure when utilizing the Berendsen barostat (Figure 14 D).

### 3.6 LLPS of Patchy Particles

In recent years, there has been growing interest in liquid-liquid phase separation (LLPS) due to the accumulation of experimental evidence showing that many cell structures are formed by this mechanism. LLPS is particularly interesting as it may explain how cells can organize themselves in a crowded environment with numerous molecular species [54]. Examples of such structures include nucleoli, Cajal bodies, stress granules, and the PSD [55].

Due to computational limitations, investigating the phase behavior of complex molecules is challenging, and coarse-graining methods are required. A minimal approach represents proteins by patchy particles, modeling multivalent interactions with attractive patches and excluded volume with repulsive interactions. [37] have used such a model to investigate the stability and composition of biomolecular condensates. PyRID is an appropriate tool for patchy particle simulations, and results from [37] has been successfully replicated for validation. See [37] for further simulation details.

Every simulation process done by PyRID is the same as in [37]. It should be noted that PyRID allows for defining binding reactions, which provides the flexibility to use a larger radius for the attractive interaction potential. Moreover, our findings suggest that the Brownian dynamics approach can reproduce the results obtained in [37] with a high level of accuracy. The coexistence curves obtained using PyRID were in agreement with [37] (Figure 15). Small deviations can probably be attributed to the slight difference in method. Whereas here, the Brownian dynamics approximation (overdamped Langevin thermostat) is utilized, [37] use a Nosé-Hoover thermostat.

**Figure 15:**
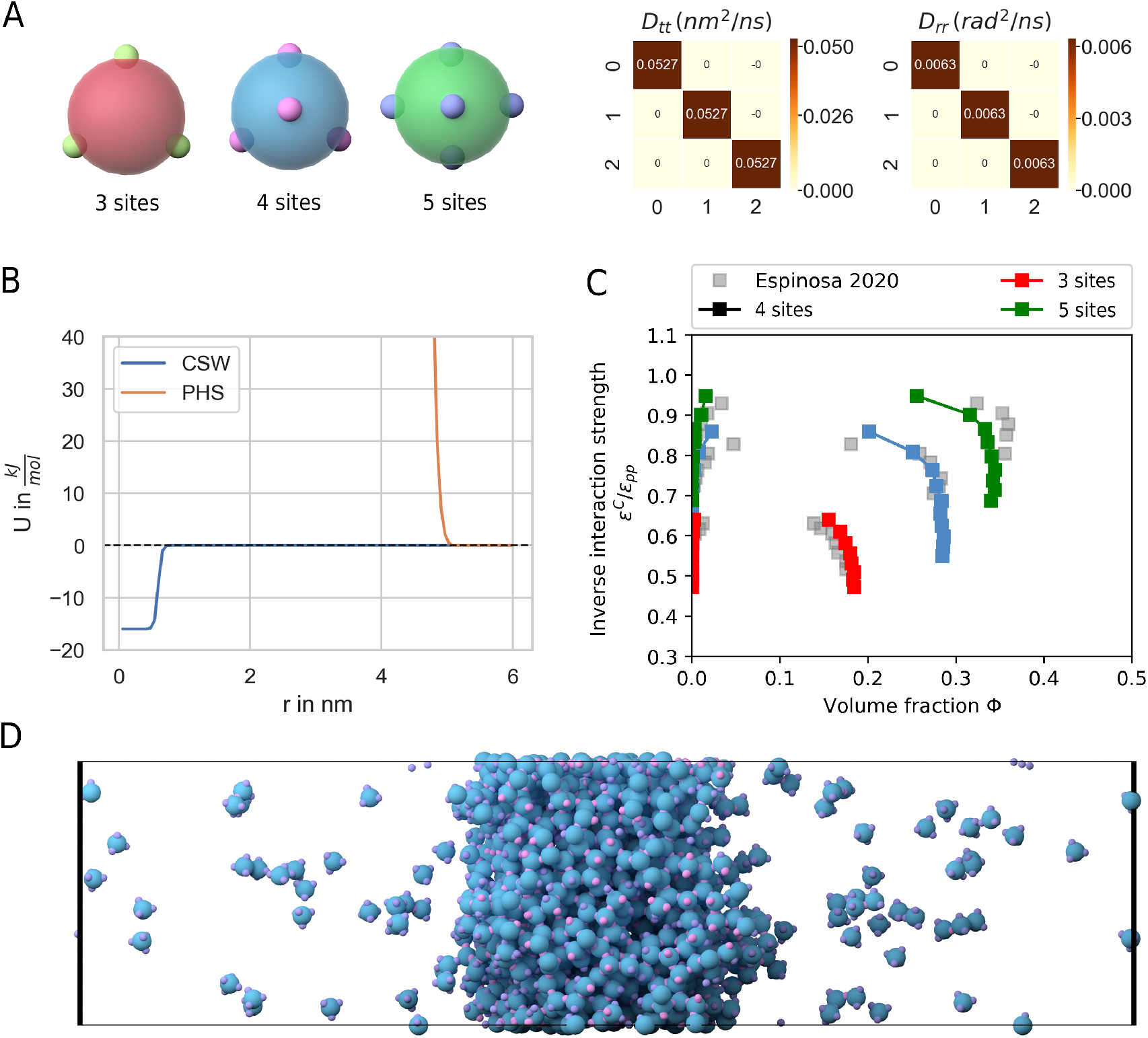
LLPS of patchy particles. (A) Left: Patchy particles with 3, 4 and 5 sites. Right: The translational and rotational diffusion tensor. (B) Graph of the continuous square-well potential (CSW) used for the attractive patches and the pseudo hard sphere potential (PHS) used for the core particle. (C) Coexistence curves for the 3, 4 and 5 sided patchy particle systems and comparison with the results from [37]. (D) Side view showing the dilute and dense phase for the 4-sided patchy particle system.

### 3.7 Benchmarks

We used a benchmark test as in ReaDDy [11], which consisted of three molecule types, A, B, and C, with radii 1.5*nm*, 3.0*nm*, and 3.12*nm*, respectively. The molecules interacted via a harmonic repulsive potential with a force constant *k* = 10*kJ/mol*, and the interaction distance *σ* was given by the radii of the interacting molecule pair:

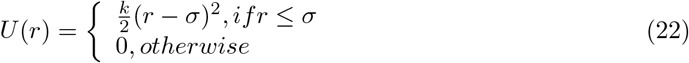

The system also included a fusion reaction 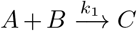 with a rate constant *k*_1_ = 0.001*ns*^*−*1^ and a reaction radius *R*_*react*_ = 4.5*nm*, as well as a fission reaction with a rate constant *k*_*−*1_ = 5.10^*−*5^*ns*^*−*1^ and a dissociation radius equal to *R*_*react*_. The benchmark was carried out for different values of the total initial molecule number *N*_*tot*_, with *NA = Ntot/4, NB = Ntot/4*, and *NB = Ntot/2*, while keeping the number density constant at *ρ*_*tot*_ = 0.00341*nm*^*−*3^. Simulations were performed for 300 ns with an integration time step of 0.1*ns*.

The performance test results showed that PyRID outperformed ReaDDy for this benchmark test (Figure 16 B, blue vs. orange curve). For particle numbers between 1,000 and 10,000, the computation time per particle update stayed approximately constant at 1.25*μs*, which corresponded to about 800,000 particle updates per second. For particle numbers above 10,000, the performance dropped slightly. The benchmark test was performed on a machine with an Intel Core i5-9300H with 2.4 GHz and 24 GB DDR4 RAM. The performance test for ReaDDy was carried out on the sequential kernel (parallel kernel were even lower in performance), and the results showed that ReaDDy scaled less linearly for large particle numbers than PyRID.

**Figure 16:**
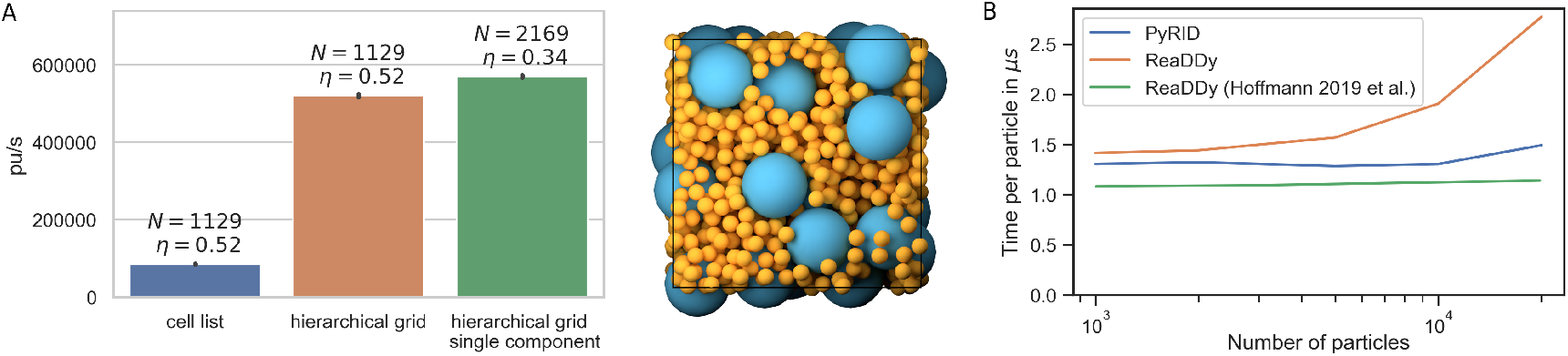
Performance test of the hierarchical grid approach. (A) Performance hierarchical grid. (B) Performance comparison between PyRID and ReaDDy. On a benchmark system with an Intel Core i5-9300H with 2.4 GHz and 24 GB DDR4 RAM, PyRID (blue line) outperforms ReaDDy (yellow). However, [11] obtained a better performance and especially scaling for ReaDDy on a different machine with an Intel Core i7 6850K processor at 3.8GHz and 32GB DDR4 RAM (green line).

The reason why ReaDDy’s scaling behavior for large particle numbers was much less linear in the benchmark test and why multi-threading led to a performance loss was not clear, but it was speculated that ReaDDy might have been compiled differently in the benchmark system used in [11] than the binaries used in this study (Figure 16 B. green curve). Nonetheless, the benchmark test showed that PyRID’s performance was comparable to ReaDDy, and in certain situations, PyRID could even outperform ReaDDy. For a system with 10^4^ particles, PyRID was able to perform at approximately 80 it/s, corresponding to approximately 7 · 10^6^ it/day. Therefore, at an integration time step of 1 ns, approximately 7 ms per day could be simulated on a medium machine.

#### Polydispersity

As mentioned in the methods chapter, PyRID uses a hierarchical grid approach to efficiently handle polydispersity. As a test, a two-component system is used. Both components consist of a single particle. Component *A* has a radius of 10 nm, component *B* has a radius of 2.5 nm. The simulation box measures 75 nm 75 nm 75 nm. The simulation volume is densely packed with both components such that we reach a volume fraction of 52%. The simulation ran for 10000 steps. When not using the hierarchical grid approach but the classical linked cell list algorithm, PyRID only reaches about 80000 particle updates per second (*pu/s*) on average (Figure 16). However, when using the hierarchical grid, more than 500000 pu/s are reached (Figure 16 A). If instead of the two component system we only simulate a one component system, PyRID also only reaches about 500000 pu/s. Thereby, PyRID performs similar independent of whether the system is mono- or polydisperse.

## 4 Discussion

In this article, we have presented PyRID, a Python-based Brownian dynamics simulator designed for interacting and reacting particles.

PyRID incorporates multiple features from existing tools, including rigid bead models for reduced particle simulation, accurate diffusion motion via experimental and theoretical diffusion tensors, and patches on bead model surfaces for multivalent protein-protein interactions. The simulator also considers polydispersity of particle sizes through a hierarchical grid data structure, supports triangulated mesh geometries to account for compartmentalized 3D environments and fixed concentration boundary conditions for simulating sub-regions within larger systems, and allows the implementation of a multitude of uni- and bimolecular reactions as well as surface diffusion. PyRID is easily modifiable and expandable by Python programmers and utilizes Numba jit compilation for efficient performance, achieving comparable results to ReaDDy.

### 4.1 Current limitations and future directions

PyRID does not incorporate hydrodynamic interactions between molecules because it would render simulations unfeasible. The calculation of the 6Nx6N diffusion tensor for the entire system is required to propagate molecule positions, and as molecule positions change at each time step, the diffusion tensor must be recalculated at each iteration. This would make large-scale simulations impractical. A discussion on this topic in the context of many-particle simulations can be found in [15]. Unlike ReaDDy, PyRID currently only supports pair interaction potentials for bond constraints, but no angular constraints for particle triplets or torsion potentials for particle quadruplets. In addition, ReaDDy uses an algorithm that ensures detailed balance for reversible reactions improving accuracy for very dense systems [56].

PyRID is a powerful tool for particle-based reaction diffusion simulations with pair-interactions, but it currently cannot simulate processes that occur over long timescales of several seconds or minutes. This limitation is significant because many cellular processes, such as signaling, self-assembly of clathrin, and protein trafficking, operate on such timescales. Other similar tools like ReaDDy [11, 50] also lack the ability to simulate on long timescales. While MCell [9] and Smoldyn [10] allow for longer simulations, they cannot resolve molecular structure or simulate protein binding and assembly. However, alternative reaction-rate based approaches that consider molecule structure, binding, and diffusion have been developed. A prominent example is NERDSS [57] that is able to resolve fast binding reactions as well as processes on large time and-spatial scales. NERDSS employs rigid body representations of molecules, like PyRID, and utilizes rejection sampling to account for excluded volume effects. However, unlike PyRID, NERDSS does not incorporate energy interaction functions, instead relying on predefined “snapping” of molecules during binding reactions to facilitate efficient simulation of assembly processes.Despite the efficiency of NERDSS in simulating assembly processes, it has limitations in predicting the structure of protein assemblies due to the predefined form of resulting assemblies. Furthermore, the lack of orientation dependence in binding reactions can lead to unrealistic events, unlike PyRID which considers orientation dependence in its construction. The use of NERDSS for describing assembly and disassembly processes relies on reaction rates measured by experimental or molecular dynamics simulation techniques. However, accurately modeling such processes using PyRID is also challenging, as it requires precise specification of energy functions governing binding interactions, which can be even more difficult to estimate than reaction rates. Additionally, NERDSS lacks interaction forces, rendering it unable to compute physical properties derived from energy functions or handle flexible chains of beads or molecules. Nonetheless, rate-based approaches like NERDSS are promising for studying complex assembly kinetics and could complement PyRID, which can enable rigid body assembly growth in principle. However, the inclusion of energy functions imposes an upper limit on the integration time step, which can be slightly increased by approximating intermolecular interaction energy functions.To address this computational bottleneck, parallelization is necessary.

For simulating very large systems, molecular dynamics tools like LAMMPS often offer parallel implementations of their algorithms using the message passing interface (MPI) standard. However, the benefits of parallelization are limited to large systems, as message passing can become a bottleneck, and may not result in significant speedup. The scalability of parallelization depends on the size of the system being simulated, and that careful consideration is required when selecting the appropriate parallelization strategy for molecular dynamics simulations.

For simulating intermediate sized systems containing 10,000-100,000 particles, GPU-based algorithms offer more promise than those designed for CPUs. A good example is HooMD [58], a molecular dynamics tool optimized for GPUs that can achieve speedups of up to two orders of magnitude compared to a single CPU and over one order of magnitude compared to a modern multi-core CPU. By harnessing the power of GPUs, particle-based reaction diffusion simulations can potentially be conducted on time scales of minutes. However, it remains unclear how efficiently the required algorithms and data structures can be ported to GPUs. Machine learning has also been integrated into simulations for various applications, such as coarse graining and molecular kinetics [59, 60]. These findings suggest that emerging technologies like GPUs and machine learning have the potential to advance simulations, but further research is needed to optimize their implementation and assess their scalability. Given that many of these tools are based on Python, we think that a Python-based simulator like PyRID can speed up a successful integration.

## 5 Acknowledgments

Supported by the German Research Foundation (DFG) via the SFB1286, projects C1, Z1.

